# Metaplasia of respiratory and digestive tissues in the Eastern oyster *Crassostrea virginica* associated with the Deepwater Horizon oil spill

**DOI:** 10.1101/2021.02.15.431248

**Authors:** Deanne S. Roopnarine, Peter D. Roopnarine, Laurie C. Anderson, Ji Hae Hwang, Swati Patel

## Abstract

Metaplasia is a well documented and deleterious effect of crude oil components on bivalved molluscs, including oysters. This reversible transformation of one cell type to another, is a common response to petroleum-product exposure in molluscs. It has been shown experimentally in previous work that eastern oysters (*Crassostrea virginica*) exposed to petroleum products will exhibit metaplasia of digestive tissues. Here we document for the first time that wild adult oysters inhabiting coastal waters in the northern Gulf of Mexico during and in the aftermath of the Deepwater Horizon oil spill (2010) exhibited metaplasia in both ctenidia and digestive epithelia at significantly higher levels than geographic controls of *C. virginica* from Chesapeake Bay. Both ctenidial (respiratory and suspension feeding) and digestive tract tissues exhibited significantly higher frequencies of metaplasia in specimens from the Gulf of Mexico compared to those from Chesapeake Bay. Metaplasia included the loss of epithelial cilia, transformations of columnar epithelia, hyperplasia and reduction of ctenidial branches, and vacuolization of digestive tissues. Evidence for a reduction of metaplasia following the oil spill (2010-2013) is suggestive but equivocal.

## Introduction

The Deepwater Horizon (DWH) oildrilling rig suffered an explosion on April 20th, 2010, resulting in loss of the rig and rupturing of the Macondo well-head on the seafloor at a depth of 1522 m. Crude oil was released into the ocean column over the next 4 months before successful shutdown of the leak. During that time it is estimated that approximately 4 million barrels of oil were released into the Gulf of Mexico (GoM) [1], making the DWH spill the world’s largest accidental spill in history. In addition, about 6.97 million liters of dispersant, a mix of surfactants and hydrocarbon solvents, were applied both at the wellhead and on surface slicks during the course of remediation efforts [2]. Sensitivity of coastal environments to spill contamination, especially salt marshes and oyster reefs, were of grave concern because of their importance to commercial fisheries, their critical role as a line of defense against coastal erosion, and because they are more difficult to ‘clean’ than barrier-island beaches [3, 4].

The immediate impacts of the spill were evident, including oil slicks, fouled beaches along states bordering the GoM, and fouled, often dead wildlife; longer-term impacts are less well understood, are still being documented, and/or remain equivocal or variable. For example, studies of the incorporation of polycyclic aromatic hydrocarbons (PAHs) into a broad array of marine organisms, including oysters, crustaceans, and fish, reported elevated levels in edible tissues through 2010, with declining levels thereafter through August 2011 [5, 6]. Small particulate and mesozooplankton assemblages collected from Mobile Bay, Alabama from August to October 2010, exhibited isotopically lighter *δ*^13^C compositions, indicative of oil incorporation into tissues, and therefore into the base of the GoM coastal food web [7], although the disturbance to community structure was transient and brief [8]. Broader changes to the food web, also during 2010, occurred as a transition from metazoan dominated benthic communities to fungal dominated assemblages [9]. Nevertheless, during the same interval, Fry and Anderson [10] recorded little change to *δ*^13^C and ^14^C content of soft-tissues in the marsh mussel *Geukensia demissa* and balanoid barnacles in the Barataria Bay estuarine region of the Mississippi River delta. This suggests either a lack or delay in the transfer of plankton-incorporated oil via food web pathways, due possibly to high standing productivity in this estuarine system [10]. Furthermore, Carmichael et al. [11] did not find evidence of oil-derived carbon or nitrogen incorporation into shell organic matrix or soft tissues of the Atlantic oyster *Crassostrea virginica* based on *δ*^13^C and *δ*^15^N values.

The extensive use of disperants in this spill, and their potential persistence in coastal areas (see [2]), also may have impacted regional ecosystems. The acute toxicity of anionic surfactants such as dioctyl sodium sulfosuccinate (DOSS) and the commercial oil-disperant products incorporating them is well documented for aquatic organisms, particularly fishes ([12] and references there in) and dispersant may have accelerated uptake of petroleum compounds by organisms exposed to dispersed oil in this spill [7]. In addition, experimental work by Bodinier et al. [12] links DOSS exposure to acute toxicity (including death) in the Gulf killifish (*Fundulus grandis*), with increased toxicity at salinities deviating the most from isosmotic values.

Ecological impacts beyond immediate and near-term mortality caused by fouling and toxic effects of oil, its components, dispersant, and dispersate-associated oil, are poorly constrained in general. Furthermore, onshore remediation efforts may have had potential ecological impacts that potentially overprinted those of the DWH spill. These efforts included large-scale diversion of Mississippi River water into Louisiana coastal wetlands during the summer of 2010 at a combined maximum flow of 780 m^3^/sec (Louisiana Office of Coastal Protection and Restoration). The diversions, particularly outflow of the Caernarvon Freshwater Diversion into Breton Sound and Davis Pond into northern Barataria Bay, altered salinity regimes and nutrient levels [13, 14], and likely the hydrodynamics of coastal marshes and nearshore waters. This sustained freshwater influx may also have had adverse effects on oysters, as the release occurred during spawning season and declines in oyster abundance, spat settlement, and filtration rates are associated with reduced salinities caused by flooding [15]. Indeed, oyster harvests on public leases were delayed east of the Mississippi River in 2010 due to a depressed spat set and oyster mortalities [16]. In addition, dramatic density decreases of oysters in areas affected by freshwater diversion efforts have been documented [17, 18], although in experimental work, Schrandt et al. [19] found that lower salinity conditions (5-10 ppt) led to increased survival of juvenile *C. virginica* exposed to oil and dispersed oil. In addition, Dietl and Durham [20] did not detect any significant difference of adult shell size between a baseline of pre-spill historical specimens and those collected in the years 2011 to 2013.

An expectation of delayed effects, and long-term consequences, of the DWH blowout is supported by studies of the Exxon Valdez spill, where both ecological and physiological impacts on a variety of marine organisms continued decades after the event [21]. An important control on any assessment of the DWH impact, however, is the establishment of the state of the GoM prior to the spill. The GoM is home to more than 4,000 offshore production platforms, with the first offshore production dating to 1937, and thousands more platforms being added in the ensuing decades [3, 22]. There have been at least five spills exceeding one million barrels prior to the DWH in the GoM, and ongoing lesser incidents and leakages (NOAA Office of Response and Restoration). There are also numerous natural, subtidal hydrocarbon seeps along the GoM continental shelf and slope (∼140,000 metric tons oil/gas per year released into the northern GoM [23]). Organisms in coastal waters of the GoM have therefore likely been exposed to naturally-seeped hydrocarbons for millenia, at least since sea level rise caused by the last glacial termination, and to hydrocarbons introduced by drilling and associated activities over the last eight decades. Additional factors complicating evaluation of spill effects include: (a) increased loads of agricultural and industrial chemicals carried into the Gulf by the Mississippi River since the 20^th^ century [24–26], and (b) fisheries methods commonly in use in the northern GOM that can profoundly alter biotic and abiotic environmental conditions [27–29].

Historical baselines of conditions within the GoM would, therefore, be useful for the proper attribution of altered conditions to DWH. Reconstructing historical baselines, however, can be limited by the availability of suitable materials, and this is particularly acute in cases where analyses depend on fresh biological materials. An alternative to reconstructing the pre-DWH state of the GoM is to substitute space for time with comparative studies of the GoM to other geographical areas among which environmental factors vary. Pennings et al. [30] used a similar approach to document recruitment failures in 2010, and subsequent recovery in 2012-14, for the marsh periwinkle *Littoraria irrorata* in Louisiana (see also [31], [32]).

The purpose of the study reported here was to examine possible impact of the DWH spill on soft-tissue morphology of the commercially significant American oyster *Crassostrea virginica*, and to monitor its continuing or diminishing effects for a period of three years after DWH, comparing specimens collected within the GoM during that time period as well as from Chesapeake Bay in Maryland, another area where *C. virginica* is subjected to anthropogenic impacts [33, 34], but without petroleum-based impacts on the scale of the Gulf of Mexico. Metaplasia, the reversible transformation of one cell type to another has been recorded previously in molluscs exposed to petroleum contamination [35–41] and, therefore, may occur in individuals exposed to DWH oil. We hypothesized that exposure to the DWH spill led to metaplasia of ctenidial (gill) and digestive tissues, although the timescale of reversal in individuals would remain unknown because of the destructive nature of individual sampling.

Evidence of DWH impact on *C. virginica* to date have been either negative or equivocal. Soniat et al. [42] recorded insignificant PAH and parasitic infection levels in tissues six months after termination of the spill, from individuals collected east of the Mississippi River in Louisiana, although there is no evidence that those specimens were directly exposed to spilled oil. Preliminary measures of heavy metal concentrations in *C. virginica* shells and tissues [43] from coastal areas of Louisiana and Alabama yielded marginally significant or insignificant values, due possibly to the low metal content of DWH oil [44] or the short residence time of some key metals in oyster tissues [45]. Likewise, linking metaplasia in other organisms to DWH has been equivocal. For example, Bentivegna et al. [46] noted a significantly higher occurrence of various lesions and metaplasia in menhaden (*Brevoortia*) from Barataria Bay compared to individuals from Delaware, but were unable to establish a significant relationship to PAH levels and the DWH spill.

Experimental exposures of bivalved molluscs to crude oil derivatives, however, have resulted in a variety of pathological responses, dependent on the contaminants (metals, hydrocarbons, etc.), including lesion development, tissue necrosis and metaplasia of gill, digestive, and renal tissues [35, 40]. Vignier et al. [47] showed that *C. virginica* larvae are capable of ingesting oil that has adhered to phytoplankton on which the larvae feed, with negative effects on larval survival, and that settled spat suffered metaplastic alterations of digestive tissues [41]. In vivo monitoring has also associated metaplasia with petroleum-based and other pollutants, including PCBs [38]. NOAA’s long-term Mussel Watch biomonitoring program [48] ranks, in order of decreasing occurrence of pathologies: metals, pesticides and PAHs [49].

The duration and frequency of metaplasia may also be indicators of the longevity of a spill’s impact. For example, the mussel *Mytilus trossulus* in Prince William Sound continued to exhibit metaplasia of the digestive gland more than five years after the Exxon Valdez spill in direct correspondence to PAH concentrations [50, 51]. Furthermore, whereas metaplasia is by definition reversible, some mollusc populations might adapt evolutionarily to chronic long-term exposure. For instance, populations of *Caryocorbula caribaea* living near active hydrocarbon seeps in Trinidad, are significantly more tolerant of PAH exposure than those living in non-seep areas [52]. Even seep-associated individuals of this species, however, exhibited pathologies when PAH concentrations were elevated experimentally above ambient seep conditions, including metaplasia of stomach epithelia. Tumour development has also been observed in *C. virginica* exposed to hydrocarbons in sediments of the northeastern United States [38].

More broadly, metaplasia may be a common molluscan response to contaminant exposure. For example, Anitha et. al [53] reported damage to the adductor muscle, mantle and gill tissues of *Crassostrea madrasensis* after exposure to copper. However, they noted that the most deleterious effects were to the epithelial tissues of the organism, noting a loss of cilia in the gills as well as vacuolization. Calabrese et. al [54] reported the same in the digestive diverticula of the blue mussel *Mytilus edulis*. Roesijadi [55] describes an overall ’emaciation’ of tissues and loss of gill structure upon exposure to metals. Berthou et al. [56] studied oyster tissue after the Amoco Cadiz oil spill in 1978 and noted changes to gonad, gill and digestive tract epithelia the year of the spill with a return to normal morphology after three to five years. The changes included atrophy and degeneration of tissues, particularly in the digestive tract.

Reversal, acclimatization, and adaptation thus all complicate the assessment of metaplasia and other tissue pathologies in response to major exposures to crude oil and petroleum products. Here we used a comparative approach to explore the occurrence and areas of metaplasia in specimens of *C. virginica* from the Gulf of Mexico which would have been exposed to the Deepwater Horizon spill, plus individuals that survived the spill, and individuals from Chesapeake Bay in Maryland which were never exposed to the spill. The major objectives of this study were to: (1) determine the types of epithelia lining the ctenidia and intestinal tract of *C. virginica* and establish whether there is evidence of metaplasia; (2) compare specimens from the Gulf of Mexico and Chesapeake Bay; (3) test whether there was a significantly elevated frequency of metaplasia in specimens from the GoM, and if so, (4) determine if the frequency varied during the time period covered by the study (2010-2013). Ctenidia and digestive tissues were selected because of their direct interaction with suspended food particles and potentially greater exposure to assimilated hydrocarbon materials. These tissues also play critical roles in the health of individuals and populations.

The ctenidia of *C. virginica* are typical lamellibranch gills, consisting of ciliated filaments, and are responsible for suspension feeding and gas exchange. Suspension feeding is accomplished by the generation of water currents and filtration of particles by arrays of ciliated epithelia lining ctenidial branch surfaces. The particular lamellibranch branching morphology, ciliary arrangements, morphologies and subsequent current system vary among lamellibranch species, but in general the ciliated linings are constructed of one to two rows of simple, columnar epithelial cells (for example, Fig. 1A). As in other ostreids, the ctenidial filaments of *C. virginica* from Chesapeake Bay are borne on prominent folded ridges (plicate ctenidia), and comprise ciliated, columnar epithelia. Ctenidial columnar cells have been observed to be underlain by cuboidal and sometimes flattened cells [57], and in the case of the oyster *Ostrea chilensis*, arise during development from ectodermal ridges on the mantle [58]. It is conventionally believed that the ctenidia serve a primarily respiratory function during larval and early post-larval developmental stages, but are clearly involved in adult feeding.

**Figure 1.**
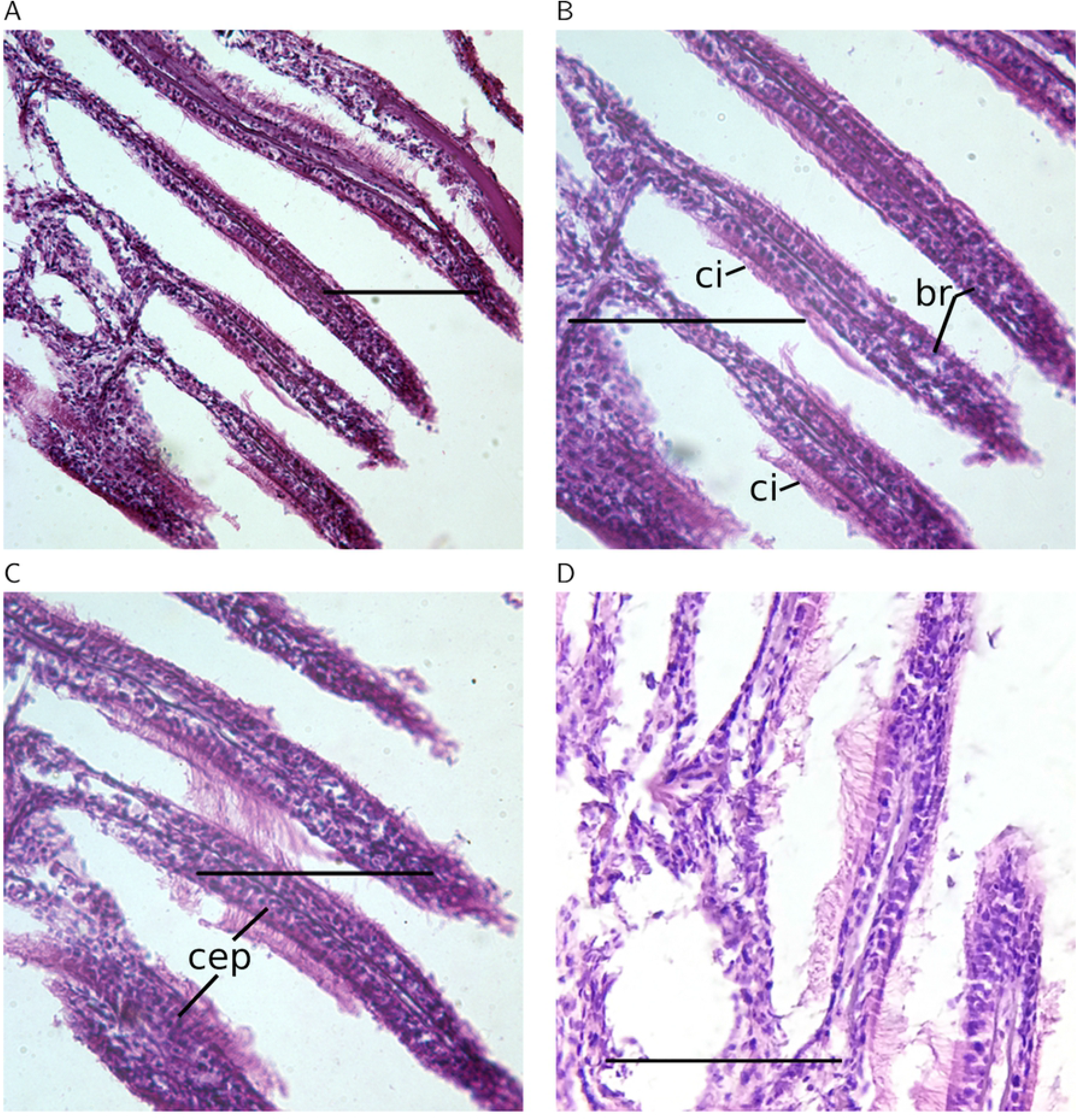
Normal ctenidial branches of *C. virginica* individuals from Chesapeake Bay. Three individuals are represented, the first in photographs A and B, and then C and D respectively. Labels: br - ctenidial branch; ci - cilia; cep - columnar epithelia. All scale bars are 100*µ*m.

We report below that the digestive system of *C. virginica* is typical of bivalves, consisting of a stomach bearing a crystalline style used in the mastication of food particles, and an upper digestive tract bearing internal, ciliated branches. Those branches presumably provide increased surface area for the absorption of digested material.

## Materials and Methods

### Specimen material

A total of 38 specimens from the Gulf of Mexico (GoM) and eight control specimens from Chesapeake Bay, Maryland, were selected for analysis. GoM specimens included those collected after initiation of the oil spill in 2010, and in subsequent years 2011 and 2013 (Tables 1 and 2). Oysters were collected in shallow subtidal areas within or immediately adjacent to *Spartina* marshes in a back-barrier lagoon connected to Barrataria Bay at the eastern end of Grand Isle, Louisiana (LA) and near the Alabama (AL) mainland along the causeway to Dauphin Island, AL in August 2010, two months after oil made landfall. Additional oysters were collected at Grand Isle in June 2013. Control specimens from Chesapeake Bay were obtained in years 2010, 2012 and 2013. Oyster clusters were collected via visual searches until at least 10 live individuals were found. Additional oysters were obtained fresh from oyster fishermen collecting in Apalachicola Bay, FL in December 2010 and December 2011. Oysters were placed on ice immediately after collection and stored below 0°C. Specimens were subsequently frozen and stored at -18° to -20°C until retrieved for analysis. Storage times varied from 3 days to 12 months.

**Table 1.**
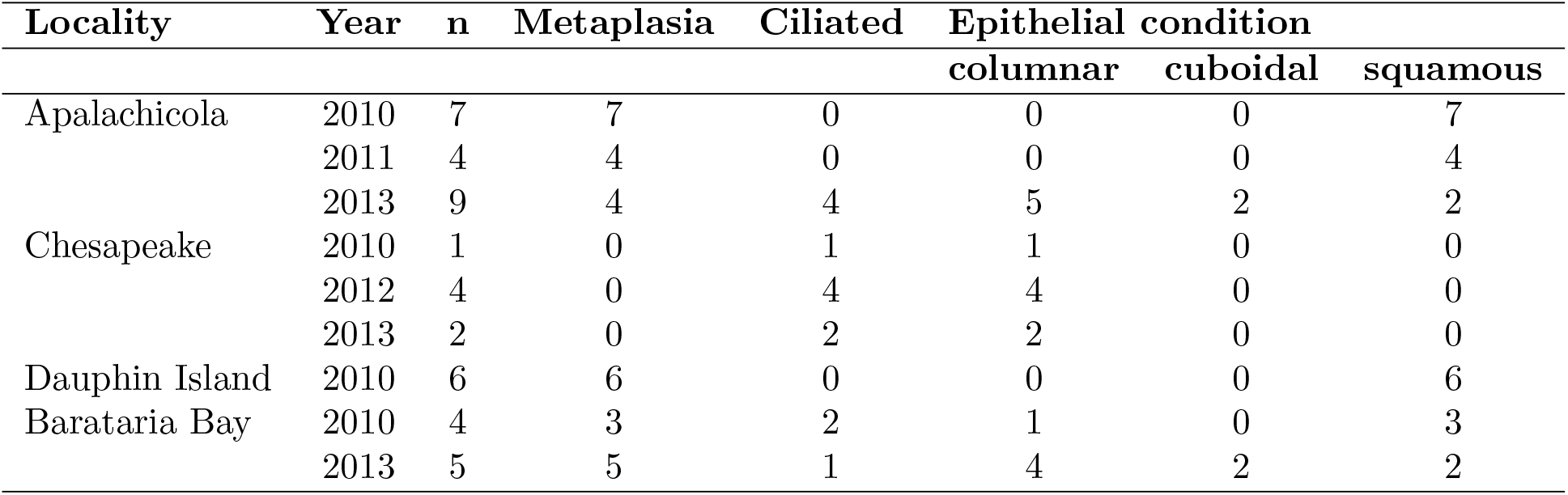
Specimens and ctenidial examinations. n is the number of specimens from each locality and year successfully examined. Ciliated - number of specimens with ciliated ctenidial branches. Epithelial condition - condition of ctenidial branch epithelia. Columnar epithelia are the normal condition, cuboidal and squamous (stratified) are metaplasial conditions. A single metaplasia specimen may possess multiple epithelial types.

| Locality | Year | n | Metaplasia | Ciliated | Epithelial condition |  |  |
| --- | --- | --- | --- | --- | --- | --- | --- |
|  |  |  |  |  | columnar | cuboidal | squamous |
| Apalachicola | 2010 | 7 | 7 | 0 | 0 | 0 | 7 |
|  | 2011 | 4 | 4 | 0 | 0 | 0 | 4 |
|  | 2013 | 9 | 4 | 4 | 5 | 2 | 2 |
| Chesapeake | 2010 | 1 | 0 | 1 | 1 | 0 | 0 |
|  | 2012 | 4 | 0 | 4 | 4 | 0 | 0 |
|  | 2013 | 2 | 0 | 2 | 2 | 0 | 0 |
| Dauphin Island | 2010 | 6 | 6 | 0 | 0 | 0 | 6 |
| Barataria Bay | 2010 | 4 | 3 | 2 | 1 | 0 | 3 |
|  | 2013 | 5 | 5 | 1 | 4 | 2 | 2 |

**Table 2.** Specimens and digestive tracts examined. n - number of specimens. Metaplasia indicates the number of specimens that exhibited some form of metaplasia, including the absence of cilia on digestive tract diverticula, and whether those diverticula were atrophied and/or possessed heavily vacuolated tissues.

| Locality | Year | n | Metaplasia | Ciliated | Diverticula |  |
| --- | --- | --- | --- | --- | --- | --- |
|  |  |  |  |  | atrophied | vacuolated |
| Apalachicola | 2011 | 2 | 2 | 0 | 2 | 2 |
|  | 2013 | 7 | 6 | 3 | 4 | 1 |
| Chesapeake | 2010 | 1 | 0 | 1 | 0 | - |
|  | 2012 | 4 | 0 | 4 | 0 | 0 |
|  | 2013 | 3 | 1 | 3 | 0 | 1 |
| Barataria Bay | 2010 | 5 | 4 | 1 | 4 | 2 |
|  | 2013 | 5 | 2 | 2 | 2 | 0 |

### Histology

Shells were opened by severing the adductor muscle as close to the right valve as possible. A macroscopic examination was then conducted of the internal surfaces of both valves and the soft tissues. Valves were examined for boring sponges, blisters, and cysts, while the soft tissue was examined for cysts and macroscopically visible parasites. Thirty eight specimens were determined to be suitable for further histological analysis, being free of macroscopic abnormalities and commensal or macro-parasitic organisms.

Gills were dissected from the body and transverse cuts made through the digestive system. Both the gills and digestive tissues were then immersed in Bouin’s fixative for 24 hours, after which they were washed in 50% alcohol for two hours and then stored in 70% ethanol for 24-48 hours. Tissue samples were subsequently dehydrated under vacuum in alcohol solutions of increasing concentration: 80% ethanol for five minutes, two changes of 95% ethanol for 10 minutes and 20 minutes respectively, and then four changes of 100% ethanol for times ranging from 15 to 30 minutes. Following dehydration, specimens were cleared in two baths of xylene, under vacuum, for 15 and 20 minutes respectively, and placed in baths for 30 minutes consisting of equal parts xylene and paraffin. Finally, specimens were infiltrated with pure paraffin from which block molds were made, and subsequently refrigerated for 24 hours at 1.6°C.

Tissue sections were sliced from each block with a rotary microtome, with sections ranging between 7-8 *µ*m in thickness. These sections were mounted on standard microscope slides with paraffin removed using two washes of xylene lasting five minutes each; tissues were re-hydrated with washes of alcohol of decreasing concentration, ranging from 100-30% for a total of five washes. Re-hydration was completed by immersion in distilled water for two minutes. Tissue sections were then stained with hematoxylin and eosin, followed again by dehydration using three washes of 100% alcohol. Slides were finally cleared with a wash of xylene prior to the application of coverslips. Slides were examined immediately after staining with a Leica DM750 microscope and images taken. More detailed examinations and microphotographs were subsequently made with a Spot Flex Ultra High Resolution CCD camera mounted on a Leica DMRB microscope.

All prepared slides are deposited in the collections of the Department of Invertebrate Zoology and Geology at the California Academy of Sciences. Shells are deposited in the collections of the South Dakota School of Mines and Technology.

## Results

A total of 46 specimens were examined, although it was not possible in all cases to examine all ctenidial and digestive tissues in the same specimen, due to the delicacy of the tissues and resulting losses during histological preparation. Both tissues were examined, however, for 18 specimens distributed among all localities (Table 1, 2). No indication of an association between the occurrence of metaplasia in ctenidial tissues and their occurrence in digestive tissues was found; we tested this observation with a binary association between the presence/absence of metaplasia in the ctenidia and digestive tract (Fisher’s exact test, *p* = 0.366). We therefore considered each tissue separately in the following analyses, testing for effects among all localities, among localities within the GoM, and among years of collection within each locality. We also checked for a dependence of the occurrence of metaplasia on the duration for which a specimen was frozen prior to histological analysis, and rejected any such dependence for either tissue (Logistic regression: ctenidia, *χ*^2^ = 0.25, *p* = 0.617; digestive, *χ*^2^ = 1.97, *p* = 0.161).

### Ctenidia

Metaplasia was manifested in ctenidia as the absence of cilia or alteration of the columnar epithelium, or both, with a significant association between the two (Fisher’s exact test, *p* = 0.0003). Twenty-one of 23 specimens lacking cilia also exhibited epithelial metaplasia, whereas nine of 13 ciliated specimens had normal, columnar epithelia (Fig. 2A). The frequency of ctenidial metaplasia varied significantly among localities, with 100% of specimens from Apalachicola Bay and Dauphin Island exhibiting some degree of alteration, 89% from Louisiana, but none from Chesapeake Bay (n = 37, Fisher’s exact test, *p* < 0.0001) (Fig. 1B; Table 1). There were no significant differences among GoM localities (n = 30, Fisher’s exact test, *p* = 0.5).

**Figure 2.**
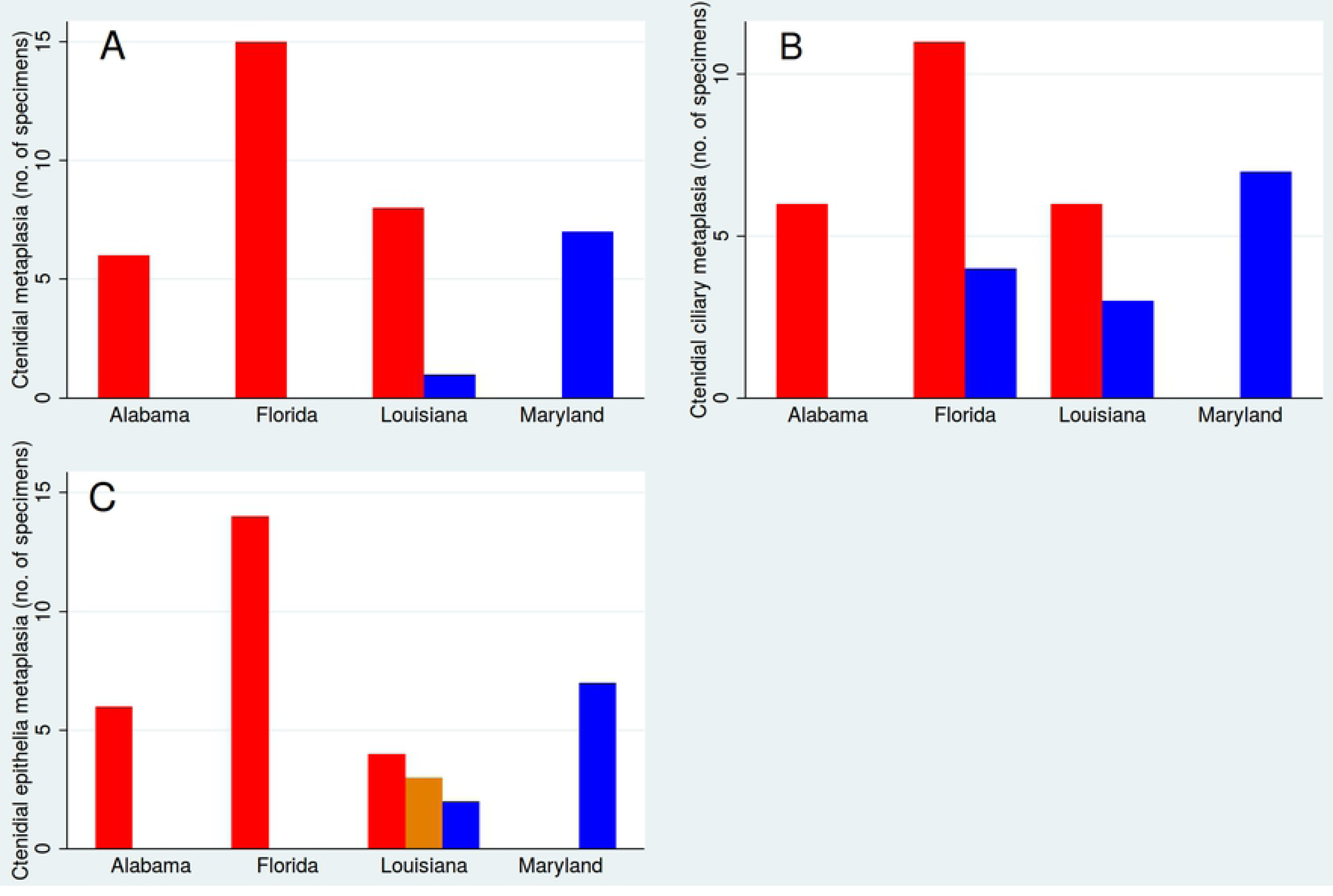
Proportions of specimens displaying ctenidial metaplasia. A - Occurrence of any type of ctenidial metaplasia (red) versus healthy individuals (blue). B - Absence of ctenidial cilia (red) versus presence (blue). C - Alteration of ctendial epithelia (red), healthy epithelia (blue), and mixed altered and healthy in the same individual (orange). Locations are Barataria Bay (Louisiana), Dauphin Island (Alabama), Apalachicola (Florida) and Chesapeake Bay (Maryland). The results are summarized for all sampled years.

Metaplasia of the cilia, defined as the absence of cilia, and exclusive of epithelial condition, varied significantly among localities solely because of the higher frequency of metaplasia in GoM samples compared to those from Chesapeake Bay (n = 37, Fisher’s exact test, *p* = 0.0011) (Fig. 2B); there was no significant difference among GoM localities when tested separately from Chesapeake Bay (n = 30, Fisher’s exact test, *p* = 0.2942) (Figure **??**). Apalachicola Bay and Louisiana samples were heterogeneous, however, having both ciliated and unciliated individuals, whereas all specimens from Dauphin Island lacked cilia. We tested for a temporal signal among collection years, comparing 2010 and 2013 for Louisiana, and 2010, 2011 and 2013 for Apalachicola Bay. There was no difference between years for Louisiana (n = 9, Fisher’s exact test, *p* = 0.5328), but significant variation among years in Apalachicola Bay (n = 15, Fisher’s exact test, *p* = 0.0015). One of 12 specimens collected from Apalachicola Bay in 2010, and another in 2011, were ciliated, whereas all three specimens collected and analyzed in 2013 were ciliated.

The presence of metaplasia in gill epithelia differed significantly among all localities (n = 36, Fisher’s exact test, *p* < 0.0001), and among GoM localities only (n = 29, Fisher’s exact test, *p* = 0.0011) (Fig. 2C). All specimens from Chesapeake Bay had the expected columnar epithelia (Fig. 1), but specimens from the GoM exhibited a variety of metaplasial alterations, such as stratified squamous epithelia (Fig. 3), and hyperplasia, a condition where the density of cells in ctenidial branches increases dramatically. Several individuals from Louisiana possessed a mixture of normal columnar and stratified squamous epithelia, and one specimen from Apalachicola Bay had mixed cuboidal and stratified squamous epithelia. Differences among the GoM localities is caused by the presence of several individuals from Louisiana with normal, columnar epithelia, whereas all individuals from Apalachicola Bay and Dauphin Island had altered epithelia. There is no evidence to support a temporal difference in Louisiana between years 2010 and 2013, however, and individuals with normal epithelia occur in both years alongside individuals with non-columnar epithelia (n = 9, Fisher’s exact test, *p* = 0.5238).

**Figure 3.**
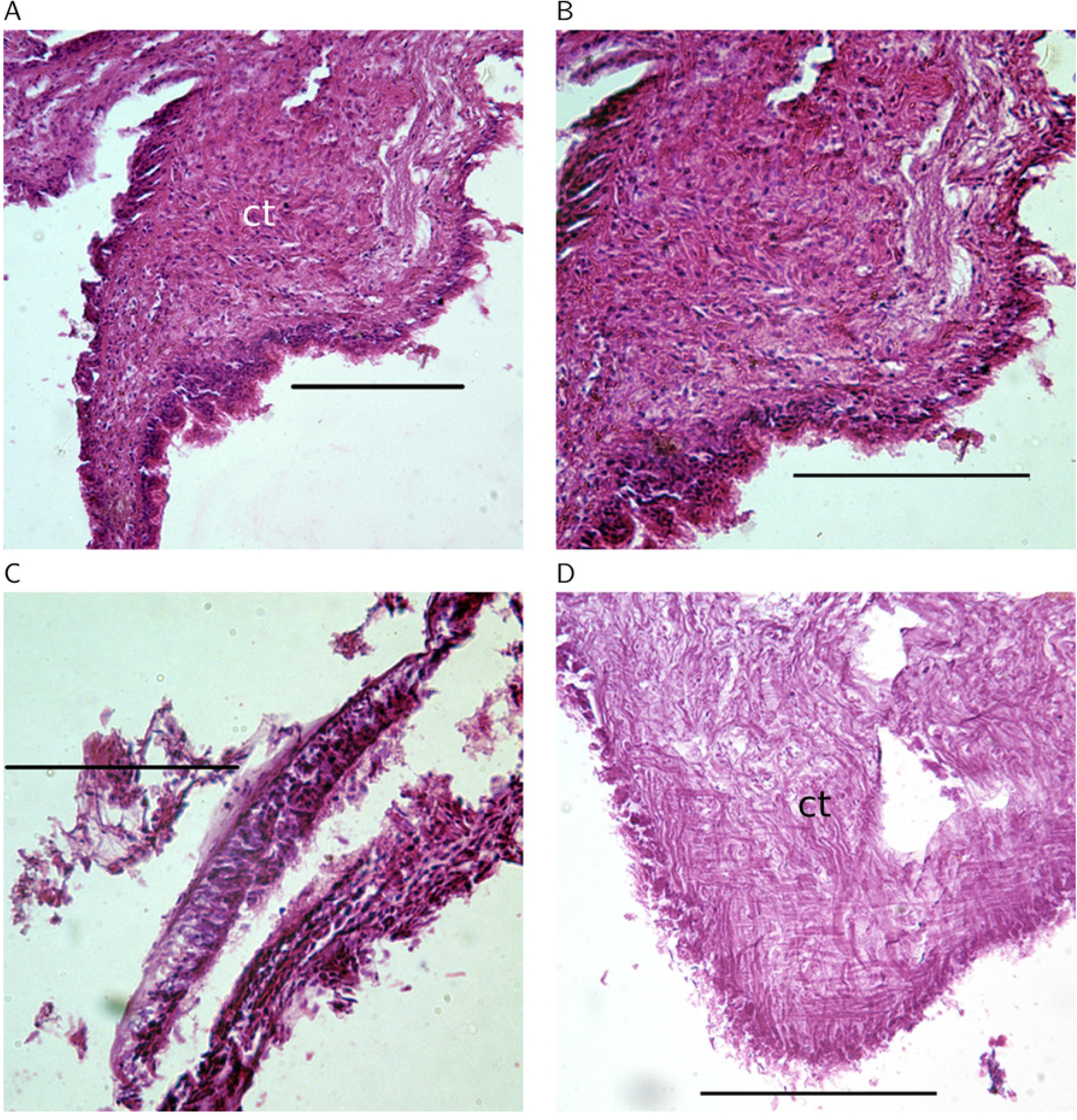
Metaplasially altered ctenidia of three individuals (A, B - Apalachicola; C - Apalachicola; D - Barataria Bay) from the Gulf of Mexico. A, B (a magnification of A) and D exhibit hyperplasia of the connective tissue and reduced epithelial structure. C exhibits stratified epithelium and a reduction of cilia on an otherwise developed ctenidial branch. Labels: ct - connective tissue. Scale bars in all images are 100*µ*m.

### Digestive tract

Three general categories of metaplasia were observed in the digestive tract: the absence of epithelial cilia, atrophy of gut diverticula, and extensively vacuolated cells (Fig. 4) (Table 2). The absence of cilia was significantly associated with atrophy of gut diverticula (Fisher’s exact test, p = 0.018), and atrophied diverticula were significantly associated with cell vacuolation (Fisher’s exact test, p = 0.010). The occurrence of any type of metaplasia differed significantly among localities (Chesapeake Bay, Apalachicola Bay, Louisiana; no digestive system tissues were successfully sectioned for specimens from Dauphin Island) (Fisher’s exact test, n = 27, p = 0.0068), and is accounted for by a significantly higher frequency in the GoM, where the two localities did not differ from each other (Fisher’s exact test, n = 18, p = 0.303) (Fig. 5). Furthermore, the difference between Chesapeake Bay and GoM localities results solely from a higher frequency of atrophied diverticula in GoM specimens (Fisher’s exact test, n = 26, p = 0.0048), and there are no significant differences with respect to the presence of cilia (Fisher’s exact test, n = 14, p = 0.293) or cell vacuolation (Fisher’s exact test, n = 17, p = 0.165). Finally, the occurrence of individuals with atrophied diverticula or vacuolated cells, in years 2010, 2011 and 2013 in Apalachicola Bay and Louisiana, renders insignificant any apparent trends in a reduction of metaplasia over that interval (Fisher’s exact test, n = 18, p = 0.354; n = 10, p = 0.080 respectively).

**Figure 4.**
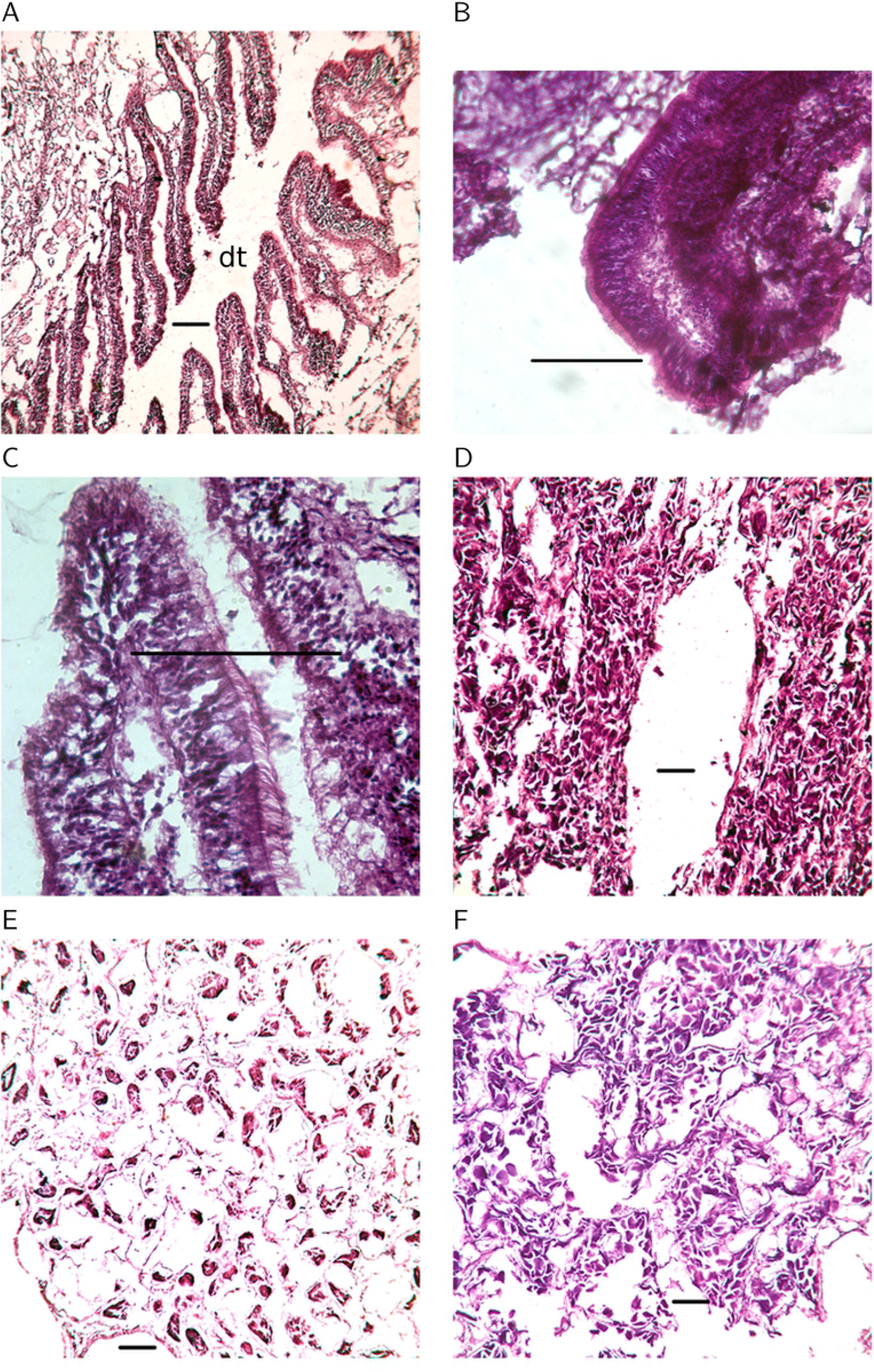
Digestive tract histologies. A-C, Normal digestive diverticula of three individuals from Chesapeake Bay. A illustrates the digestive tract (dt) and diverticular branches, and B and C exhibit ciliated columnar epithelia lining the diverticula. exhibiting ciliated columnar epithelia. D-F, three individuals from Barataria Bay, Louisiana, displaying atrophy and vacuolization of tissue resulting in breakdown of epithelial integrity. D - digestive tract with atrophied diverticula. E - epithelium along the digestive tract showing a loss of epithelial integrity. F - A digestive tract consisting of multiple atrophied tubules. Scale bars in all images are 100*µ*m.

**Figure 5.**
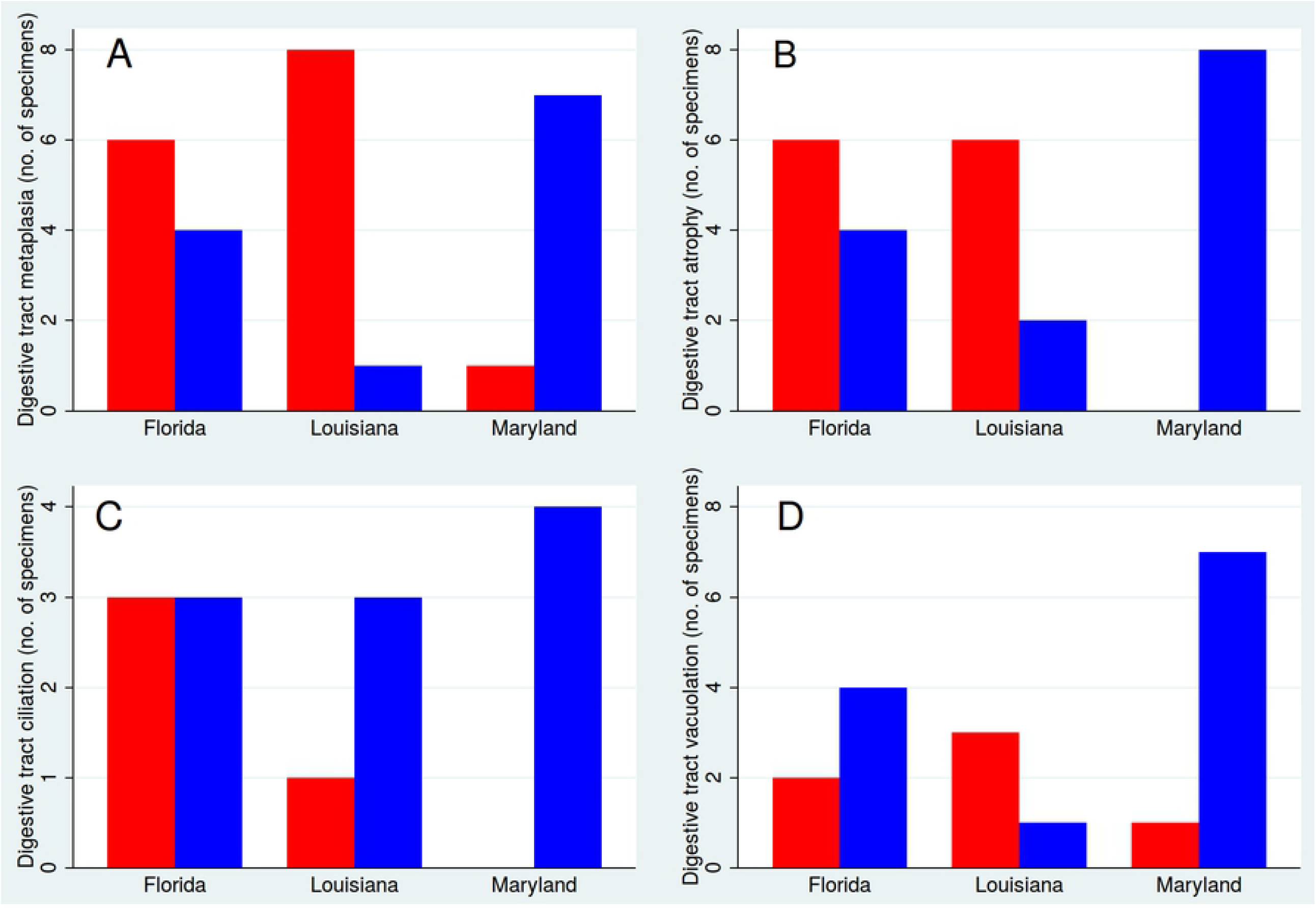
Proportions of specimens displaying digestive tract metaplasia. A - Occurrence of any type of digestive metaplasia (red) versus healthy individuals (blue). B - Atrophy of tract diverticula (red) versus normal diverticula (blue). C - Absence of digestive tract cilia (red), versus ciliated tracts (blue). D - Extensive cellular vacuolation (red), versus non-vacuolated cells (blue). Locations are Barataria Bay (Louisiana), Apalachicola (Florida) and Chesapeake Bay (Maryland).

## Discussion

We found that specimens of *Crassostrea virginica* from Chesapeake Bay display ctenidial and digestive tissue morphologies and histologies consistent with those described previously for molluscan bivalves in general, and ostreid bivalves specifically. The frequency of ctenidial metaplasia was significantly greater in the Gulf of Mexico than in Chesapeake Bay during the interval covered by this study, but no significant differences among localities within the GoM were found. Ctenidial metaplasia included the absence of epithelial cilia and alteration of the expected columnar epithelium to cuboidal or squamous cells, or a combination of all those cell types. A lack of cilia was always associated with some type of epithelial alteration, but epithelial cell type alteration could occur without a corresponding lack of cilia. Furthermore, variation in ciliary metaplasia is indicated; all specimens from Alabama were unciliated, collections from Louisiana always contained some ciliated individuals, regardless of the year in which they were collected, whereas collections from Florida showed a significantly declining incidence of metaplasia during the study, that is, an increasing frequency of healthy, ciliated individuals. Whether this indicates a reversal of the conditions that induced ciliary metaplasia in the first place would have to be determined on the basis of continued monitoring.

Metaplasial variation of the ctenidial epithelium was complex both within and between individuals from the GoM, but some form of metaplasia was present in specimens from all localities. Both the cuboidal and squamous alterations represent a decrease in the height of the epithelia compared to normal columnar cells, and in most cases represented a partial to complete collapse of ctenidial branches. A qualitative conclusion drawn from our examination of the altered tissues is that both the absence of columnar cells, replaced instead by cuboidal or squamous cells, and the morphology of the altered branches, result in tissues that bear a strong resemblance to the ectodermal ridges from which the ctenidial structure develops ontogenetically. We therefore speculate that metaplasia in this case could consist of, or include a reversion to an ontogenetically earlier developmental morphology. Furthermore, in contrast to the ciliary condition, there is no indication of temporal variation or a reversal of metaplasia in the epithelial tissues during the sampling interval.

Both the digestive tract and stomach exhibited significant metaplasia in specimens from the Gulf of Mexico, whereas specimens from Chesapeake Bay differed little or not at all from the expected morphologies. There was no clear association of metaplasia between the ctenidia and digestive system in our specimens, however; metaplasia of ctenidial tissues was not predictive of metaplasia of digestive tissues and vice versa. The lack of association points to a complex relationship between the stressors that induce metaplasia and the ways in which individuals may respond.

Digestive tissues exhibited a similar array of metaplasial conditions, comprising an absence of cilia, atrophy of the digestive tract, and vacuolation of gastric tissues. Specimens from the GoM displayed a significantly higher frequency of metaplasia compared to those from Chesapeake Bay, and this was due solely to the occurrence of digestive tract atrophy in the GoM, a condition which was completely absent in Chesapeake Bay specimens. Additionally, the absence of cilia and tissue vacuolation, although both significantly associated with digestive tract atrophy, were also present in some specimens from Chesapeake Bay. Finally, there was no evidence of a declining incidence of metaplasia in GoM specimens during the sampling interval (2010-2013).

Metaplasial tissues may be expected to perform physiologically differently from unaltered tissues, and their presence has implications for individual and population performance [59]. An overall implication, therefore, of metaplasia of respiratory and digestive tissues is a decreased physiological performance of individuals. For example, cilia on ctenidial branches are responsible for particle capture and sorting during filter feeding, moving food particles toward the labial palps for subsequent ingestion. The ctenidia themselves feature significantly in gas exchange, and squamous or cuboidal epithelia form ctenidia with less surface area for efficient gas exchange in comparison to those constructed from columnar epithelia. Similarly, atrophied digestive tracts have reduced surface area for the release of digestive enzymes and the subsequent absorption of nutrients. It is also possible that the composition of the suite of digestive enzymes secreted during the digestive process is altered as a consequence of atrophy. A reduction or absence of cilia within the digestive tract would result in reduced movement of ingested materials through the tract, and possibly hinder egestion. We speculate that the vacuolation of digestive tissues has a negative impact on digestive and absorptive surface area, as well as the production and secretion of enzymes. It has been established that diseased, stressed or non-feeding bivalves, including *C. virginica*, possess metaplasial digestive tract epithelia [59–61].

The frequency of metaplasia documented in this study would suggest first, that specimens from the GoM perform more poorly in comparison to those from Chesapeake Bay, and second that performance perhaps varied during the sampling interval of the study, with respiratory performance improving over time. If these results can be extrapolated to the populations which were sampled, then one would expect a decline of population productivity in the GoM between 2010 and 2013. This expectation is based both on physiological performance, as well as aspects of population dynamics, such as reproductive output, because of the involvement of physiological systems in reproduction. For example, although species of *Crassostrea* are broadcast spawners, the ctenidial and inhalant-exhalant water systems are involved in the internal movement and spawning of fertilized gametes [62].

An outstanding question that motivated this study is if the Deepwater Horizon (DWH) oil spill, and mitigation measures such as dispersants, can be implicated in the types and frequencies of metaplasia observed [41]. Evidence in support of such a suggestion are the significantly higher frequencies observed in the GoM compared to Chesapeake Bay, and perhaps the observed reduction of ctenidial metaplasia during the interval following the spill. If this is indeed the case, it should be noted that the specimens used for this study were not directly contaminated with DWH oil, even the August 2010 Grand Isle, Louisiana specimens. Therefore the impacts of spills may not have to result from direct contact with a fresh, undiluted slick and can still be measured months to years after initial contamination [21, 63–68].

There are, however, several difficulties with establishing support for this hypothesis. First, no baseline exists for the occurrence and types of metaplasia in the areas sampled. Whereas it is feasible that variation of metaplasial type and frequencies are the result of variations of exposure to DWH oil, it is equally likely given current evidence that metaplasia within the GoM is the result of either exposure to naturally occurring hydrocarbons (from natural seeps), or that it is a now common feature within the GoM because of the long history of petroleum exploration, production and contamination there. An alternative hypothesis is that our inference of metaplasia in the GoM oysters is incorrect, and the tissue anomalies are in fact the result of evolutionary change driven by long-term exposure of the GoM population to both naturally occurring and anthropogenically derived hydrocarbons. A similar explanation was offered for the occurrence of abnormal tissues in bivalves sampled from within oil fields off Trinidad [52]. This explanation could be consistent with the fact that our specimens from Chesapeake Bay do not exhibit the abnormalities so common in the GoM, and would therefore be the result of much greater exposure to hydrocarbons in the GoM, both historically and because of the Deepwater Horizon spill. Alternatively, given the reportedly great sensitivity of oysters in the Chesapeake system to chronic exposure to pollutants, it remains unclear why our specimens from that population are significantly healthier than specimens from the GoM, unless the former is a positive benefit of ongoing mitigation efforts in the Chesapeake system. Resolving these questions requires more spatially and temporally extensive sampling and monitoring both within geographic areas of concern, and areas considered to have fewer or distinct anthropogenic impacts.

## Acknowledgments

Field work in Louisiana and Alabama was assisted by Annette Engel, Carrol Michael, and Caroline Dietz. Timothy Chung, Michael Hellman and Scott Elliott assisted with histological preparations. The authors acknowledge support from The Nova Southeastern University President’s Faculty Research and Development Grant No. 335317, and field support from the Louisiana Sea Grant College Program.

## References

1. McNutt MK, Camilli R, Crone TJ, Guthrie GD, Hsieh PA, et al. (2012) Review of flow rate estimates of the Deepwater Horizon oil spill. Proceedings of the National Academy of Sciences 109: 20260–20267.

2. White HK, Lyons SL, Harrison SJ, Findley DM, Liu Y, et al. (2014) Long-term persistence of dispersants following the Deepwater Horizon oil spill. Environmental Science & Technology Letters 1: 295–299.

3. Davis D, Baumann R (2001) Oil in Louisiana’s estuarine environments: A development model. Science in China Series B: Chemistry 44: 230–239.

4. Fleeger J, Riggio M, Mendelssohn I, Lin Q, Deis D, et al. (2019) What promotes the recovery of salt marsh infauna after oil spills? Estuaries and Coasts 42: 204–217.

5. Fernando H, Ju H, Kakumanu R, Bhopale KK, Croisant S, et al. (2019) Distribution of petrogenic polycyclic aromatic hydrocarbons (PAHs) in seafood following Deepwater Horizon oil spill. Marine Pollution Bulletin 145: 200–207.

6. Xia K, Hagood G, Childers C, Atkins J, Rogers B, et al. (2012) Polycyclic aromatic hydrocarbons (PAHs) in Mississippi seafood from areas affected by the Deepwater Horizon oil spill. Environmental Science & Technology 46: 5310–5318.

7. Graham WM, Condon RH, Carmichael RH, D’Ambra I, Patterson HK, et al. (2010) Oil carbon entered the coastal planktonic food web during the Deepwater Horizon oil spill. Environmental Research Letters 5: 045301.

8. Carassou L, Hernandez F, Graham W (2014) Change and recovery of coastal mesozooplankton community structure during the Deepwater Horizon oil spill. Environmental Research Letters 9: 124003.

9. Bik HM, Halanych KM, Sharma J, Thomas WK (2012) Dramatic shifts in benthic microbial eukaryote communities following the Deepwater Horizon oil spill. PLoS One 7: e38550.

10. Fry B, Anderson LC (2014) Minimal incorporation of Deepwater Horizon oil by estuarine filter feeders. Marine pollution bulletin 80: 282–287.

11. Carmichael RH, Jones AL, Patterson HK, Walton WC, Pèrez-Huerta A, et al. (2012) Assimilation of oil-derived elements by oysters due to the deepwater horizon oil spill. Environmental Science & Technology 46: 12787–12795.

12. Bodinier C, Galvez F, Gautreaux K, Rivera A, Green CC (2015) Toxicological and physiological effects of the surfactant dioctyl sodium sulfosuccinate at varying salinities during larval development of the gulf killifish (Fundulus grandis). In: Impacts of Oil Spill Disasters on Marine Habitats and Fisheries in North America, CRC Press Taylor & Francis Group. pp. 35–51.

13. Bianchi TS, Cook RL, Perdue EM, Kolic PE, Green N, et al. (2011) Temperature impacts of diverted freshwater on dissolved organic matter and microbial communities in Barataria Bay, Louisiana, U.S.A. Marine Environmental Research 72: 248–257.

14. McDonald T, Telander A, Marcy P, Oehrig J, Geggel A, et al. (2015) Temperature and salinity estimation in estuaries of the northern gulf of mexico. NOAA Tech Rep https://wwwfwsgov/doiddata/dwh-ar-documents/863/DWH-AR0270936pdf.

15. Pollack JB, Kim HC, Morgan EK, Montagna PA (2011) Role of flood disturbance in natural oyster (Crassostrea virginica) population maintenance in an estuary in South Texas, USA. Estuaries and Coasts 34: 187–197.

16. Tunnell Jr JW, Feinberg KR (2011) An expert opinion of when the Gulf of Mexico will return to pre-spill harvest status following the BP Deepwater Horizon MC 252 oil spill. Gulf Coast Claims Facility, Feinberg Rozen, LLP.

17. Grabowski JH, Powers SP, Roman H, Rouhani S (2017) Potential impacts of the 2010 Deepwater Horizon oil spill on subtidal oysters in the Gulf of Mexico. Marine Ecology Progress Series 576: 163–174.

18. Powers SP, Grabowski JH, Roman H, Geggel A, Shahrokh R, et al. (2017) Consequences of large-scale salinity alteration during the Deepwater Horizon oil spill on sutidal oyster populations. Marine Ecology Progress Series 576: 175–187.

19. Schrandt J, Powers S, Rikard FS, Thongda W, Peatman E (2018) Short-term low salinity mitigates effects of oil and dispersant on juvenile eastern oysters: A laboratory experiment with implications for oil spill response activities. PLOS ONE 13: e0203485.

20. Dietl GP, Durham SR (2016) Geohistorical records indicate no impact of the Deepwater Horizon oil spill on oyster body size. Royal Society Open Science 3: 160763.

21. Peterson CH, Rice SD, Short JW, Esler D, Bodkin JL, et al. (2003) Long-term ecosystem response to the Exxon Valdez oil spill. Science 302: 2082–2086.

22. Priest T (2007) Extraction not creation: the history of offshore petroleum in the Gulf of Mexico. Enterprise & Society 8: 227–267.

23. Kennicutt MC (2017) Oil and gas seeps in the Gulf of Mexico. In: Habitats and Biota of the Gulf of Mexico: Before the Deepwater Horizon Oil Spill, Springer. pp. 275–358.

24. Rabalais NN, Turner RE, Sen Gupta BK, Platon E, Parsons ML (2007) Sediments tell the history of eutrophication and hypoxia in the northern Gulf of Mexico. Ecological Applications 17: S129–S143.

25. Trefry JH, Metz S, Trocine RP, Nelsen TA (1985) A decline in lead transport by the Mississippi River. Science 230: 439–441.

26. Turner R, Rabalais N, Justic D (2006) Predicting summer hypoxia in the northern Gulf of Mexico: Riverine N, P, and Si loading. Marine Pollution Bulletin 52: 139–148.

27. Beck MW, Brumbaugh RD, Airoldi L, Carranza A, Coen LD, et al. (2011) Oyster reefs at risk and recommendations for conservation, restoration, and management. Bioscience 61: 107–116.

28. Lenihan HS, Peterson CH (1998) How habitat degradation through fishery disturbance enhances impacts of hypoxia on oyster reefs. Ecological Applications 8: 128–140.

29. Queirós A, Hiddink JG, Kaiser M, Hinz H (2006) Effects of chronic bottom trawling disturbance on benthic biomass, production and size spectra in different habitats. Journal of Experimental Marine Biology and Ecology 335: 91–103.

30. Pennings SC, Zengel S, Oehrig J, Alber M, Bishop TD, et al. (2016) Marine ecoregion and Deepwater Horizon oil spill affect recruitment and population structure of a salt marsh snail. Ecosphere 7: e01588.

31. Zengel S, Montague CL, Pennings SC, Powers SP, Steinhoff M, et al. (2016) Impacts of the deepwater horizon oil spill on salt marsh periwinkles (Littoraria irrorata). Environmental Science & Technology 50: 643–652.

32. Zengel S, Weaver J, Pennings SC, Silliman B, Deis DR, et al. (2017) Five years of Deepwater Horizon oil spill effects on marsh periwinkles Littoraria irrorata. Marine Ecology Progress Series 576: 135–144.

33. Black H, Andrus CFT, Lambert W, Rick T, Gillikin D (2017) δ^15^N values in Crassostrea virginica shells provides early direct evidence for nitrogen loading to Chesapeake Bay. Scientific Reports 7: 44241.

34. Kemp WM, Boynton WR, Adolf JE, Boesch DF, Boicourt WC, et al. (2005) Eutrophication of Chesapeake Bay: historical trends and ecological interactions. Marine Ecology Progress Series 303: 1–29.

35. Neff JM, Hillman RE, Carr RS, Buhl RL, Lahey JI (1987) Histopathologic and biochemical responses in Arctic marine bivalve molluscs exposed to experimentally spilled oil. Arctic: 220–229.

36. Guzmán-García X, Botello A, Martinez-Tabche L, González-Márquez H (2009) Effects of heavy metals on the oyster (Crassostrea virginica) at Mandinga Lagoon, Veracruz, Mexico. Revista de Biología Tropical 57: 955–962.

37. Sindermann C (1982) Implications of oil pollution in production of disease in marine organisms. Philosophical Transactions of the Royal Society of London B, Biological Sciences 297: 385–399.

38. Gardner GR, Yevich PP, Harshbarger JC, Malcolm AR (1991) Carcinogenicity of Black Rock Harbor sediment to the eastern oyster and trophic transfer of Black Rock Harbor carcinogens from the blue mussel to the winter flounder. Environmental Health Perspectives 90: 53–66.

39. Osman G, Galal M, Abul-Ezz A, Mohammed A, Abul-Ela M, et al. (2017) Polycyclic aromatic hydrocarbons (PAHS) accumulation and histopathological biomarkers in gills and mantle of Lanistes carinatus (Mollusca, Ampullariidae) to assess crude oil toxicity. Punjab University Journal of Zoology 32: 39–50.

40. Gregory M, Marshall D, George R, Anandraj A, McClurg T (2002) Correlations between metal uptake in the soft tissue of perna perna and gill filament pathology after exposure to mercury. Marine Pollution Bulletin 45: 114–125.

41. Vignier J, Rolton A, Soudant P, Fu-lin EC, Robert R, et al. (2018) Evaluation of toxicity of Deepwater Horizon slick oil on spat of the oyster Crassostrea virginica. Environmental Science and Pollution Research 25: 1176–1190.

42. Soniat TM, King SM, Tarr MA, Thorne MA (2011) Chemical and physiological measures on oysters (Crassostrea virginica) from oil-exposed sites in Louisiana. Journal of Shellfish Research 30: 713–717.

43. Roopnarine P, Roopnarine D, Gillikin DP, Anderson LC, Ballester M, et al. (2011) Uptake of heavy metals and pahs from the deepwater horizon oil spill by soft tissues and shells of the coastal oyster crassostrea virginica. In: AGU Fall Meeting Abstracts. volume 2011, pp. B33E–0517.

44. Alcazar F, Kennicutt II MC, Brooks JM (1989) Benthic tars in the Gulf of Mexico: Chemistry and sources. Organic Geochemistry 14: 433–439.

45. Thompson KF, Kennicutt II MC, Brooks JM (1990) Classification of offshore Gulf of Mexico oils and gas condensates (1). AAPG Bulletin 74: 187–198.

46. Bentivegna CS, Cooper KR, Olson G, Pena EA, Millemann DR, et al. (2015) Chemical and histological comparisons between Brevoortia sp.(menhaden) collected in fall 2010 from Barataria Bay, LA and Delaware Bay, NJ following the DeepWater Horizon (DWH) oil spill. Marine Environmental Research 112: 21–34.

47. Interactions between Crassostrea virginica larvae and Deepwater Horizon oil: Toxic effects via dietary exposure, author=Vignier, J and Rolton, A and Soudant, Philippe and Chu, FLE and Robert, Rene and Volety, AK, journal=Environmental Pollution, volume=246, pages=544–551, year=2019, publisher=Elsevier.

48. Kimbrough KL, Lauenstein G, Christensen J, Apeti D (2008) An assessment of two decades of contaminant monitoring in the Nation’s Coastal Zone. NOAA/National Centers for Coastal Ocean Science.

49. Kim Y, Powell EN, Wade TL, Presley BJ (2008) Relationship of parasites and pathologies to contaminant body burden in sentinel bivalves: NOAA Status and Trends ‘Mussel Watch’ Program. Marine Environmental Research 65: 101–127.

50. Carls MG, Heintz R, Moles A, Rice SD, Short JW (2001) Long-term biological damage: What is known, and how should that influence decisions on response, assessment, and restoration? In: International Oil Spill Conference. American Petroleum Institute, volume 2001, pp. 399–403.

51. Babcock M, Harris P, Carls M, Brodersen C, Rice S (1998) Mussel Bed Restoration and Monitoring, Exxon Valdez Oil Spill Restoration Final Project Report (Restoration Project 95090). National Oceanic and Atmospheric Administration, National Marine Fisheries Service, Auke Bay Laboratory, Juneau, Alaska.

52. Mohammed A, Agard J (2004) The occurrence of NADPH-ferrihemoprotein reductase in Corbula caribea, from a natural oil seep at La Brea, Trinidad. Marine Pollution Bulletin 48: 784–789.

53. Anitha P, George RM (2006) Histopathology of copper toxicity in the Indian edible oyster, Crassostrea madrasensis (Preston). Journal of the Marine Biological Association of India 48: 19–23.

54. Calabrese A, MacInnes JR, Nelson DA, Greig RA, Yevich PP (1984) Effects of long-term exposure to silver or copper on growth, bioaccumulation and histopathology in the blue mussel Mytilus edulis. Marine Environmental Research 11: 253–274.

55. Roesijadi G (1996) Metallothionein and its role in toxic metal regulation. Comparative Biochemistry and Physiology part C: Pharmacology, Toxicology and Endocrinology 113: 117–123.

56. Berthou F, Balouet G, Bodennec G, Marchand M (1987) The occurrence of hydrocarbons and histopathological abnormalities in oysters for seven years following the wreck of the Amoco Cadiz in Brittany (France). Marine Environmental Research 23: 103–133.

57. Elston R (1980) Functional anatomy, histology and ultrastructure of the soft tissues of the larval American oyster, Crassostrea virginica. In: Proceedings National Shellfisheries Association.

58. Chaparro O, Videla J, Thompson R (2001) Gill morphogenesis in the oyster Ostrea chilensis. Marine Biology 138: 199–207.

59. Lowe DM, Moore MN, Clarke KR (1981) Effects of oil on digestive cells in mussels: quantitative alterations in cellular and lysosomal structure. Aquatic Toxicology 1: 213–226.

60. Winstead JT (1995) Digestive tubule atrophy in eastern oysters, Crassotrea virginica (Gmelin, 1791), exposed to salinity and starvation stress. Journal of Shellfish Research 14: 105–112.

61. Eble AF, Scro R (1996) General anatomy. The Eastern oyster Crassostrea virginica Maryland Sea Grant College, Maryland: 19–73.

62. Foighil DÓ, Taylor DJ (2000) Evolution of parental care and ovulation behavior in oysters. Molecular Phylogenetics and Evolution 15: 301–313.

63. Vandermeulen J (1982) Some conclusions regarding long-term biological effects of some major oil spills. Philosophical Transactions of the Royal Society of London B, Biological Sciences 297: 335–351.

64. Vandermeulen J, Singh J (1994) Arrow oil spill, 1970–90: Persistence of 20-yr weathered bunker C fuel oil. Canadian Journal of Fisheries and Aquatic Sciences 51: 845–855.

65. Reddy CM, Eglinton TI, Hounshell A, White HK, Xu L, et al. (2002) The West Falmouth oil spill after thirty years: the persistence of petroleum hydrocarbons in marsh sediments. Environmental Science & Technology 36: 4754–4760.

66. Effect of field exposure to 38-year-old residual petroleum hydrocarbons on growth, condition index, and filtration rate of the ribbed mussel, Geukensia demissa, author=Culbertson, Jennifer B and Valiela, Ivan and Olsen, Ylva S and Reddy, Christopher M, journal=Environmental Pollution, volume=154, number=2, pages=312–319, year=2008, publisher=Elsevier.

67. Esler D, Iverson SA (2010) Female harlequin duck winter survival 11 to 14 years after the Exxon Valdez oil spill. The Journal of Wildlife Management 74: 471–478.

68. Li H, Boufadel MC (2010) Long-term persistence of oil from the Exxon Valdez spill in two-layer beaches. Nature Geoscience 3: 96.

